# “Patchiness” in Mechanical Stiffness across a Tumor as an Early-Stage Marker for Malignancy

**DOI:** 10.1101/2023.07.31.551398

**Authors:** Zibah Mirzakhel, Gudur Ashrith Reddy, Jennifer Boman, Brianna Manns, Savannah Van Teer, Parag Katira

## Abstract

Mechanical phenotyping of tumors, either at an individual cell level or tumor cell population level is gaining traction as a diagnostic tool. However, the extent of diagnostic and prognostic information that can be gained through these measurements is still unclear. In this work, we focus on the heterogeneity in mechanical properties of cells obtained from a single source such as a tissue or tumor as a potential novel biomarker. We believe that this heterogeneity is a conventionally overlooked source of information in mechanical phenotyping data. We use mechanics-based in-silico models of cell-cell interactions and cell population dynamics within 3D environments to probe how heterogeneity in cell mechanics drives tissue and tumor dynamics. Our simulations show that the initial heterogeneity in the mechanical properties of individual cells and the arrangement of these heterogenous sub-populations within the environment can dictate overall cell population dynamics and cause a shift towards the growth of malignant cell phenotypes within healthy tissue environments. The overall heterogeneity in the cellular mechanotype and their spatial distributions is quantified by a “patchiness” index, which is the ratio of the global to local heterogeneity in cell populations. We observe that there exists a threshold value of the patchiness index beyond which an overall healthy cell population of cells will show a steady shift towards a more malignant phenotype. Based on these results, we propose that the “patchiness” of a tumor or tissue sample, can be an early indicator for malignant transformation and cancer occurrence in benign tumors or healthy tissues. Additionally, we suggest that tissue patchiness, measured either by biochemical or biophysical markers, can become an important metric in predicting tissue health and disease likelihood just as landscape patchiness is an important metric in ecology.

## Introduction

Cancer arises from malfunctioning cells where a deregulation of normal signaling pathways leads to the acquisition of hallmark features such as chronic proliferation and evasion of apoptosis, which disrupts normal tissue structure and function (1–4). While the majority of focus in literature has been on how changes to biological signaling pathways can aid cancerous behavior, less attention has been drawn to the effects of altering mechanical signaling pathways (2–4). Modifications to these mechanical signaling pathways can disrupt a variety of cellular processes such as cell division, cell death, and mechanotransduction - a process by which cells sense their physical environment and relay mechanical cues into biochemical signals (5,6). Malignant transformation has been associated with changes to both extra-cellular and intra-cellular mechanical properties, with studies suggesting that these altered physical properties are significant in promoting tumor progression (2,4,7–9). One such manifestation of these altered mechanical properties in malignant cells is that they are significantly softer than their healthy counterparts, an observation consistent across many cancer types (4,10–12). This decrease in cell stiffness has been linked to certain cancerous features like uncontrollable proliferation, evasion of apoptosis, and increase in motility (3,11,13).

Studies have suggested that the stiffness of cells can grade its metastatic potential, where highly invasive malignant cells are on an average (at the population level) softer than less invasive cells (14,15). Since no two individuals are perfectly identical, cell stiffness measurements, irrespective of stiffness measurement techniques, produce a distribution of cell stiffness values for any given cell population. This distribution arises from phenotypic heterogeneity across individual cells and can be persistent over several cell generations (16–18) (Figure 1). While the heterogeneity in cell mechanical properties (such as cell stiffness or adhesion) has been noticed and reported previously, it is the mean values of these distributions that is used to differentiate between the two cell types, and potentially grade the malignancy and metastatic potential of a particular cell population (9,17–19). We posit that, the distribution in cell mechanical properties across a population of cells comprising a tumor or even a normal tissue can provide significant insight into how the population will evolve over time and lead potential malignancy and metastasis (figure 1). This position is based on the fact that even healthy cell populations have broad distributions in cell mechanical properties indicating the presence of at least a small number of cells that have cancer-type phenotypes at the extremes of these distributions (14,16–18,20,21). Additionally, recent work on micron-scale, in-situ mechanical characterization of tumors has shown that tumors are made up of regions of soft and stiff cells (22), and the definitions and organization of these regions correlates with the aggressiveness of tumors (23). We build on this position with the help of computational simulations of cell-cell interactions within a 3D tissue environment, observing the evolution of cell population with heterogenous mechanical properties over time. We focus mainly on the cell stiffness as a key mechanical property where individual cells show heterogeneity, and specifically look for the growth of cells with lower stiffness (softer cells) since this is a cell trait which is strongly associated with cancer (16,17,24).

**Figure 1.**
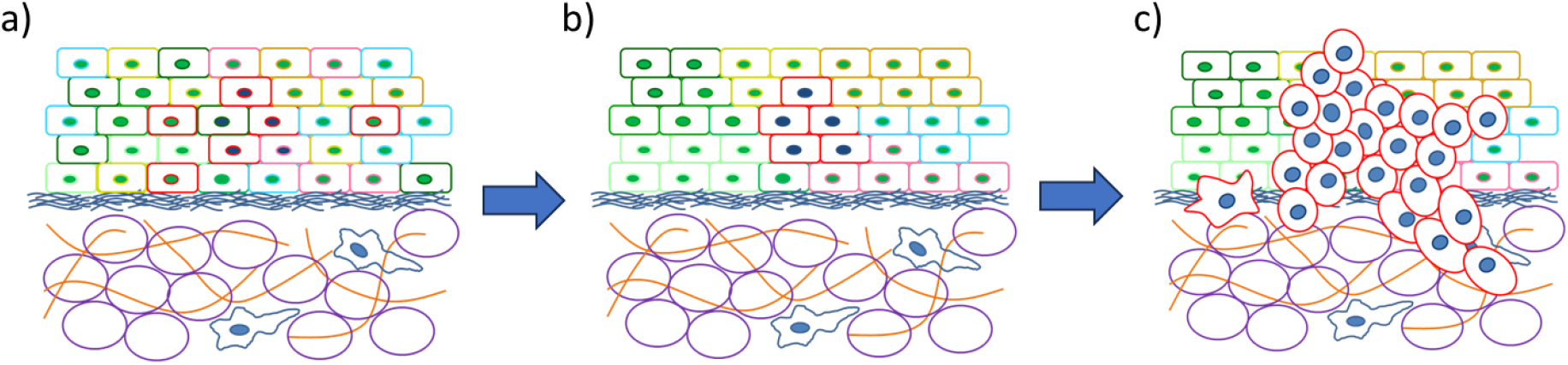
a) Mechanical phenotypic diversity in a healthy tissue, forming a spatially homogeneous tissue b) Mechanically similar cells are in close proximity with one another, forming a spatially heterogeneous tissue, c) Based on spatial arrangement and mechanical variation within the tissue, some cells are able to replicate at a faster rate, overtaking the tissue and migrating through the basement membrane.

## Methods

Mechanics-based mathematical and computational models have been a common tool used to isolate and study the sole effect that specific tumor mechanical properties have on cancer progression (2). Many multi-scale models exist, all intended to answer specific questions regarding cancer mechanics and tumor progression (3,25–27). We have previously developed one such model to understand how a mechanically distinct populations of stiff and soft cells can interpenetrate(28,29) or promote and drive tumor growth within stiff environments (3,29). The focus in these prior works was on the interactions between two mechanically distinct phenotypes of cells. Here, we build on these models to incorporate a heterogeneous population of cells with stiffness values distributed around a predetermined mean corresponding to a healthy cell’s stiffness.

In the model, cells are described as viscoelastic shells, characterized by a dense actin cortex and a liquid core, able to compete for space while interacting with other cells in the tissue (3) (Figure 2). The position of each cell is defined by a single point while the cell shape and its neighbors are obtained by Voronoi Tessellations about these individual cell points. The polyhedral cell area, volume and the interface between neighboring cells are used to compute the mechanical energy stored within each cell at any given time using Equation 1 (3,28)-

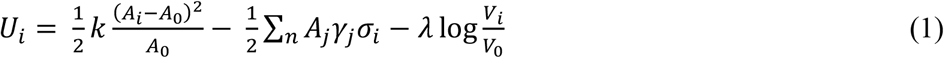

**Figure 2.**
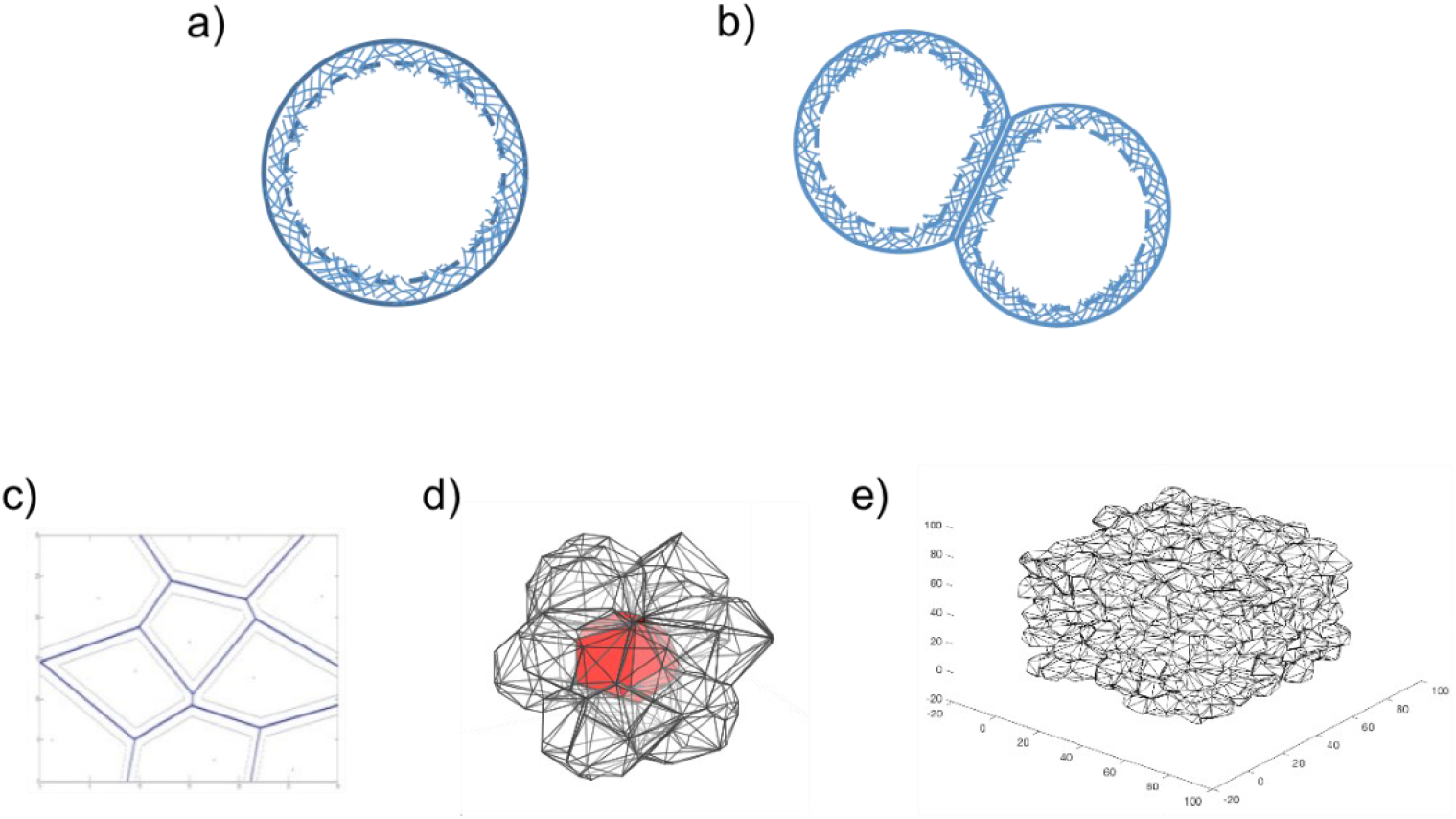
a) Individual cells are modeled as spherical in free solution, b) cells interact with one another by forming a flat interface with neighboring cells, c) a 2D representation of cells interacting with one another in tissue, where cells are modeled as Voronoi polyhedrons, d) a 3D representation of a cluster of cells modeled as Voronoi polyhedrons, e) a tissue comprised entirely of cells

The equation consists of three terms which estimate the energy of the stretched actin shell, the energy released by formation of inter-cellular bonds and the work done by the osmotic pressure inside the cell in changing its volume from *V*_0_ in free solution to *V*_*i*_ inside the tissue. *k* is the stiffness of the actin cortex, *A*_*i*_ is the area of the cell, and *A*_0_ is the area of the cell had it been spherical, n is the number of neighbors, and *A*_*j*_, *σ*_*i*_, and *γ*_*j*_ are the area of the interface, the bond density, and the bond energy between the cell and its j^th^ neighbor, respectively. *λ* is calculated as *P*_*0*_*V*_*0*_, where *P*_*0*_ and *V*_*0*_ are the osmotic pressure and volume of a cell free in solution. Cellular rearrangements, obtained by displacing the individual cell points, are dependent on the overall mechanical energy of the system (sum of mechanical energies of individual cells). Rearrangements that lower the total mechanical energy of the system (∑*U*_*i*_) are always accepted, while rearrangements that increase the energy of the system are accepted based on the probability of acceptance given by equation 2, following the Monte-Carlo Metropolis algorithm (3).

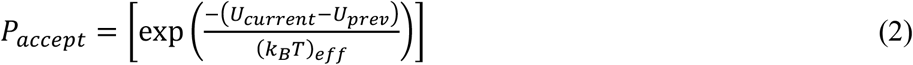

The term U_current_ refers to the tissue’s total energy in the current configuration, U_prev_ refers to the total energy in the previous configuration and (k_b_T)_eff_ refers to the internal energy of the cells, analogous to the work a cell can do via filopodial protrusions (3).

Cell fate (death or division) in the model system is a stochastic function of cell stretch, in accordance with what has been observed experimentally (30,31), and as given by equations 3 and 4 (3).

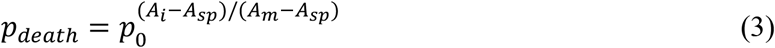

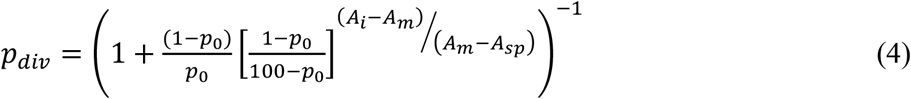

Cell death and division events drastically increase the system energy and are thus interspersed between a large number (∼70,000) of cellular rearrangements that drive the tissue back to a lower energy state as dictated by (*k*_*B*_*T*)_*eff*_. The parameter values used in the above calculations are summarized in supplementary table 1.

The initial tissue/tumor configuration for a homogeneous tissue environment is obtained by randomly dispersing all the cell defining points into a 3D space (Figure 2e). These location points act as the centers of their encapsulating, polyhedral cells. A log-normal distribution of cell stiffness values with a fixed mean of 500 Pa and different amounts of variance is specified for each cell to model mechanical heterogeneity in the tissue (17,24,32). Either high, moderate, or low amounts of mechanical heterogeneity is considered, modeled by changing the mode of the log-normal distribution. The primary results presented here use log-normal distributions for cell stiffness values to relate most closely with experimental measurements (17,21). The mode of the log-normal distributions for the different populations are set to either 465 Pa, 455 Pa, 445 Pa, 435 Pa or 425 Pa (corresponding to a standard deviation of 115 Pa, 130 Pa, 145 Pa, 160 Pa or 175 Pa respectively).

In addition to modeling the mechanical variation, the effects of the spatial arrangement of cells within the healthy tissue are also tested, as cell-to-cell interaction may further impact tumor incidence (3) (Figure 3). To setup up the initial conditions for a spatially heterogeneous tissue environment, a small number of cells (8, 27, or 64) are first randomly distributed into the 3D space and their mechanical properties are randomly chosen from the overall population distribution. The rest of the tissue is then sequentially seeded by placing cells with randomly assigned mechanical stiffness values in proximate locations to existing mechanically similar cells. This leads to clustering of mechanically similar cells while maintaining overall population heterogeneity. By altering the number of seed cells first introduced, we can alter the sizes of the clusters in the tissue (Figure 3). The size of the resulting clusters is dependent on the number of initial seed cells, where fewer seed cells amount to larger, but fewer clusters of cells with similar mechanical properties in the healthy tissue. For each spatial arrangement and stiffness distribution, a complementary non-clustered tissue is simulated as a control. A large number of cell rearrangements are performed immediately after seeding all the cells (∼70,000), without any cell death or division, in order to reach an initial low energy configuration for the model tissue system.

**Figure 3.**
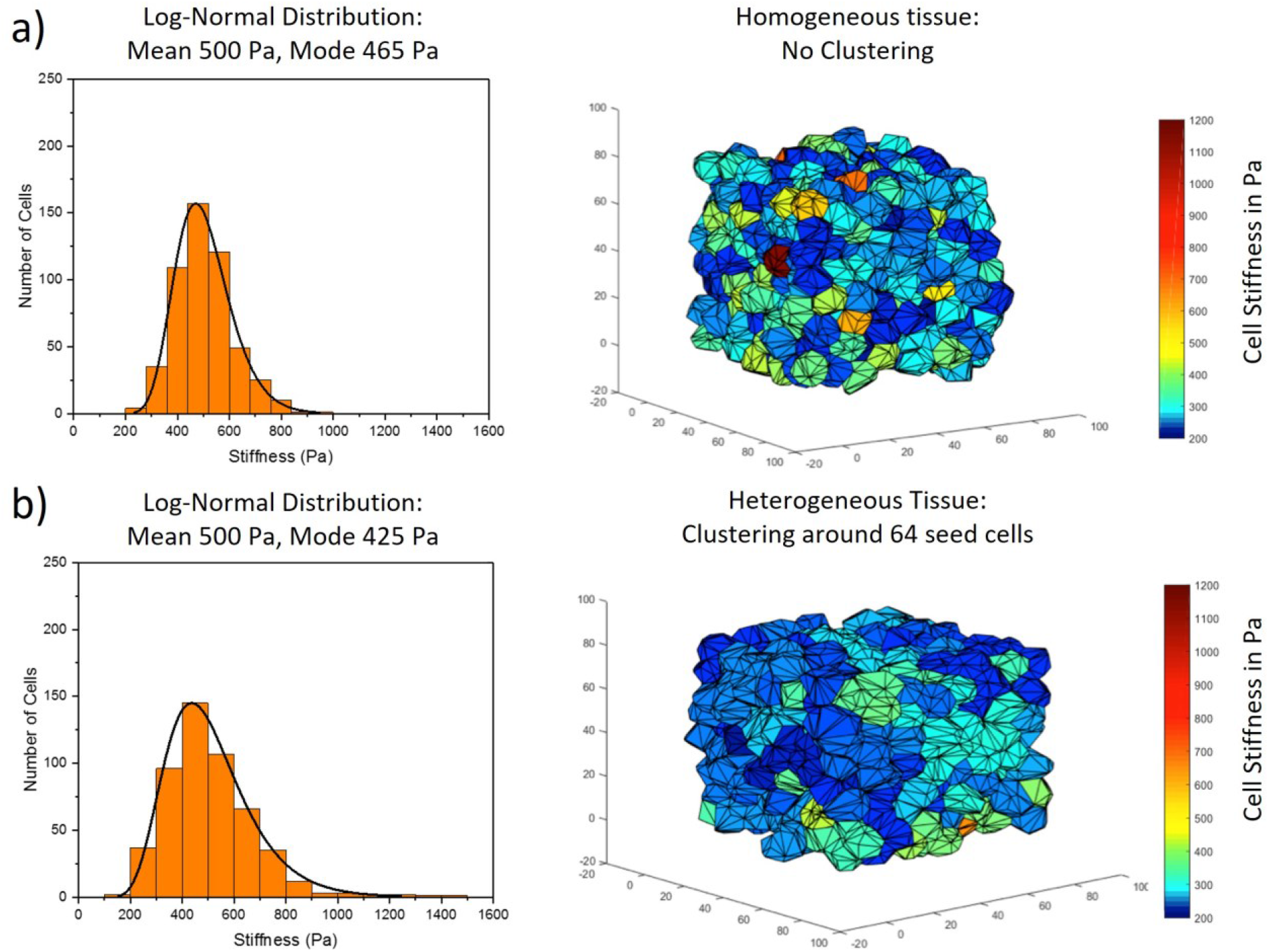
Initial healthy cell stiffness distributions applied to tissue with various spatial arrangements a) Log-normal cell stiffness distribution with a mean of 500 Pa and mode 465 Pa (standard deviation of 115 Pa) applied to a spatially homogeneous tissue b) Log-normal distribution cell stiffness distribution with a mean of 500 Pa and mode 425 Pa (standard deviation of 175 Pa) applied to a spatially non-homogeneous tissue, where clusters are formed between mechanically similar cells (64 seed cells around which other cells are sequentially populated).

This simulation adequately mimics the environment of a dense 3D tissue or tumor where cells are packed together in packing fractions close to 1. Cellular rearrangements drive the tissue to a steady energetic state. When a tissue is in an energetically steady state, cell fate decisions are made based on the stochastic equations 3 and 4, where cell stretch and area provide a driving signal for cell death and division (30). Empirically, perfectly spherical cells have a low probability of cell division and a high probability of cell death (33), while more stretched out cells with larger areas have a low probability of death and a high probability of division (34). By using this probabilistic method of cell death and division, an important aspect of cell fate decisions – mechanosensing, is integrated within the model while preserving the homeostatic nature of the tissue. When the cell divides, the resulting two daughter cells will either retain the same mechanical stiffness as its parent cell or vary slightly, where mechanical phenotype is strongly conserved (35–37). We assume a ±5% (max) variation in daughter cell stiffness compared to the parent cell. After cell fate decisions are made, cellular rearrangements again drive the tissue to a steady state energy value with the same Monte-Carlo Metropolis thermalization process. Cell rearrangement takes place over a timescale of the order of days, whereas the cell death and division processes occur within a couple of hours (38). This separation of timescales allows for the cellular rearrangement and cell death and division events to be simulated sequentially.

We simulate an 80 µm x 80 µm x 80 µm cubic space, containing 256 cells (average cell volume of ∼ 1000 µm^3^) with periodic boundary conditions. Here we assume that healthy cell population has an average cell stiffness value of 500 Pa (starting mean stiffness of all cells in the tissue). Individual cells within this population with a stiffness value lower than 400 Pa are considered tumor-like, based on experimental results that determine that cancer cells are found to be at least 20% softer than healthy cells (19,24,32).

The changes in the cell populations are tracked over 20 cellular rearrangement and death/division cycles, to determine if there are any shifts within the population and if there is an increase in the tumor-like cell population. For analysis purposes, we assume that a drop of 15 Pa in the mean cell stiffness within the tissue environment over the 20 death and division cycles implies a malignant transformation of the tissue. This drop in mean tissue stiffness can be a result of an increase in the softer cell population, a decrease in the stiffer cell population, or a combination of both. We can track the number of healthy (stiffness > 400 Pa) and cancer cells (stiffness < 400 Pa) within the population. The choices of tissue size, cell numbers, and simulation cycles are primarily restricted by the computational cost of this model. However, we believe the overall insight from this model system can be applied to larger collection of cells in tissues.

To quantify global vs local heterogeneity within the tissue environment, we borrow the concept of patchiness from environmental ecology (39–42). The patchiness index is calculated as the ratio of global to local heterogeneity using equation 5,

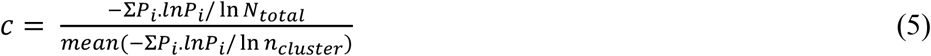

where, *P*_*i*_ is the fraction of the population within the sample that is similar (in this case has similar cell stiffness) and *N* (*N*_*total*_ or *n*_*cluster*_) is the population size of the sample. The numerator is the global heterogeneity calculated for the total population, while the denominator is the local heterogeneity calculated for a small region of the system. To calculate the *P*_*i*_, the cell stiffness values are combined into 10 Pa increment bins, and P_i_ is calculated as the ratio of the number of cells in bin *i* to the total number of cells. The same process is used for both local and global populations. The local population heterogeneity is obtained using a moving window approach, where a cubic window of predetermined dimensions is sampled, the heterogeneity of the cell population within that window is calculated, and the window is then shifted by specified distance to obtain a new local sample population. The mean of the local heterogeneities thus calculated is used in equation 5 to obtain the patchiness index. Here we use window sizes of 30 µm x 30 µm x 30 µm, 40 µm x 40 µm x 40 µm, and 50 µm x 50 µm x 50 µm and shift the window by ∼ 1 cell length (10 µm) each time, along each of the three dimensions.

## Results

As mentioned above, we use a decrease in mean cell stiffness by 15 Pa over 20 simulation cycles as the benchmark value when considering tumor occurrence and malignant transformation. This value is determined to be significant based on the constant drop of the mean value of the distribution, as well as the increase of tumor-like cells in the tissue after every cell fate decision. If the simulated tissue were to undergo more cell death and division cycles, one would expect larger drops in the mean value as well as an increased number of tumor-like cells in the tissue (Supplementary Figure 1). In the instances when tumor incidence is observed as defined by a 15 Pa drop in mean cell stiffness, it is often accompanied by an increase in tumor-like cells and a drop in the number of stiffer cells (see supplementary figure 2). However, there are also instances when the number of tumor-like cells in the tissue remains constant and the number of stiffer cells drops, which we believe is due to the inherent stochasticity in our system (see supplementary figure 3). In rare cases, the mean value of the distribution increases by 15 Pa and is accompanied by increases in the number of stiffer cells and a drop in the number of tumor-like cells (see supplementary figure 4). Although this is infrequent, it may be beneficial to understand the interplay between the mechanical variation and spatial clustering, and how it can sustain the growth of stiffer cells.

Overall, we simulate tissue systems with 5 different initial cell stiffness distributions (log-normal distributions with mean 500 Pa and modes - 465 Pa, 455 Pa, 445 Pa, 435 Pa or 425 Pa). For each of these 5 tissue systems, we further simulate four different levels of clustering within the tissue (high – 8 seed cells, medium – 27 seed cells, low – 64 seed cells and no clustering). For each of these 20 scenarios (listed in supplementary table 2), we simulate 10-12 unique instances of cell population evolution using our model, running each system for 20 cell death/division and reorganization cycles. For each of the main 20 scenarios, we count the runs where malignant transformation is observed as defined by a decrease in the mean stiffness by 15 Pa and divide it by the total number of simulations run for that scenario to find the probability of malignant transformation. We observe that both the mechanical variance and spatial arrangement collectively play influential roles in increasing the likelihood of malignant transformation (Figure 4). Individually, these factors are not significant enough in inducing tumorigenesis but are instrumental in doing so together. As both the mechanical variation and clustering of cells with similar mechanical stiffness increases in an initial healthy tissue, the probability for tumor occurrence increases (see Figure 4).

**Figure 4.**
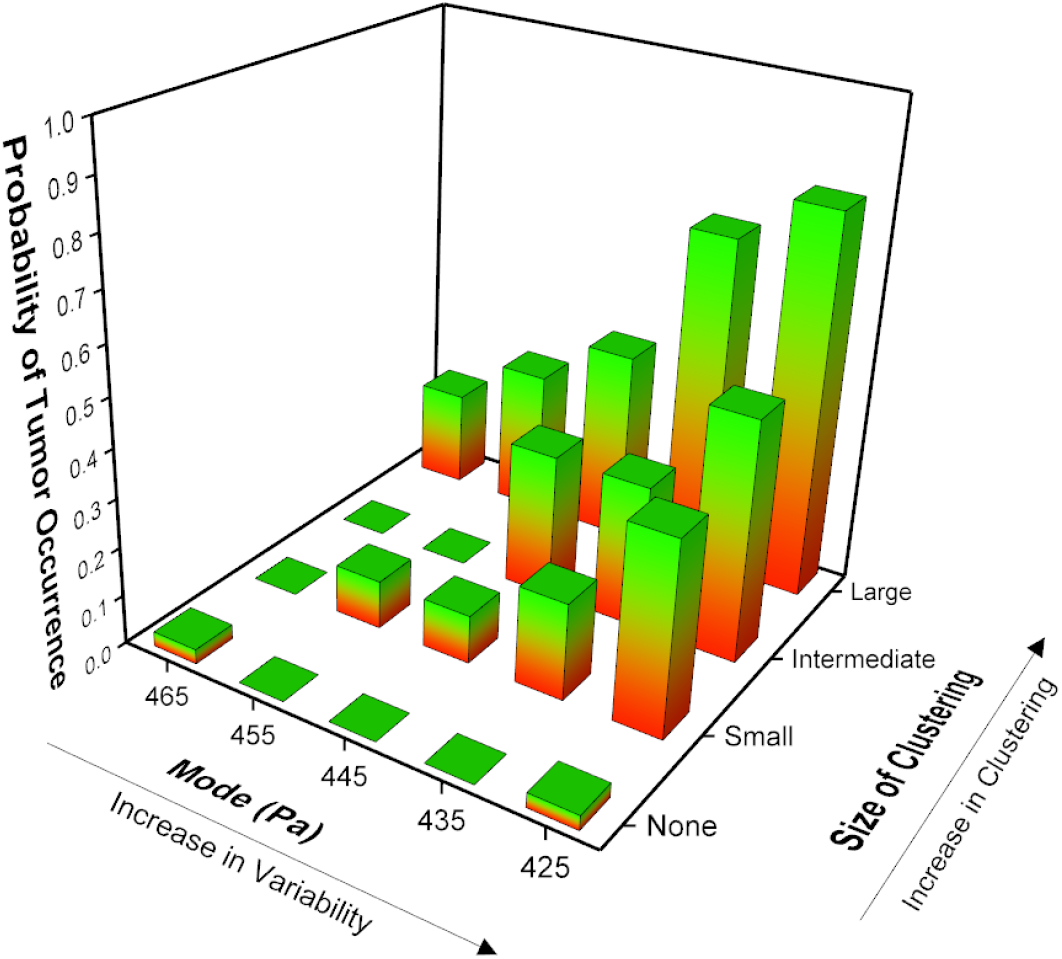
The probability of tumor occurrence is dependent on increases in the mechanical variance and the spatial clustering between cells with similar mechanical properties.

The need for spatial clustering of cells with similar mechanical properties for tumorigenesis reveals the dependency of interactions between tumor-like cells that is needed to promote their increased proliferation. This may be due to the enlarged surface area of the tumor-like cell clusters within the tissue, which gives them the ability to resist the force imposed by their surrounding stiff cells (3). This can be related to what has been seen experimentally, where tumor cells are able to withstand their stiff environments as they grow and multiply, driven by an increased homeostatic pressure for these cells (13,43).

We combine the effect of overall variance (heterogeneity) in cell mechanotype and the local clustering of mechanically similar cells using the patchiness index for the tissue. We find that independent of the cubic window size used to estimate the patchiness index (3 cell lengths, 4 cell lengths or 5 cell lengths along each dimension), there is a significant likelihood that tissues with patchiness index greater than 0.85 to show a decrease in mean cell stiffness and thus a transition towards a malignant mechanotype (figure 5, supplementary figure 5. In these figures, the stars denote p<0.05 on a single population ttest, indicating a non-zero change in mean cell stiffness for tissues with that patchiness index. We note here that we are limited in our analysis to these small window sizes because of the small size of our simulation domain (8 cell lengths along each dimension).

**Figure 5.**
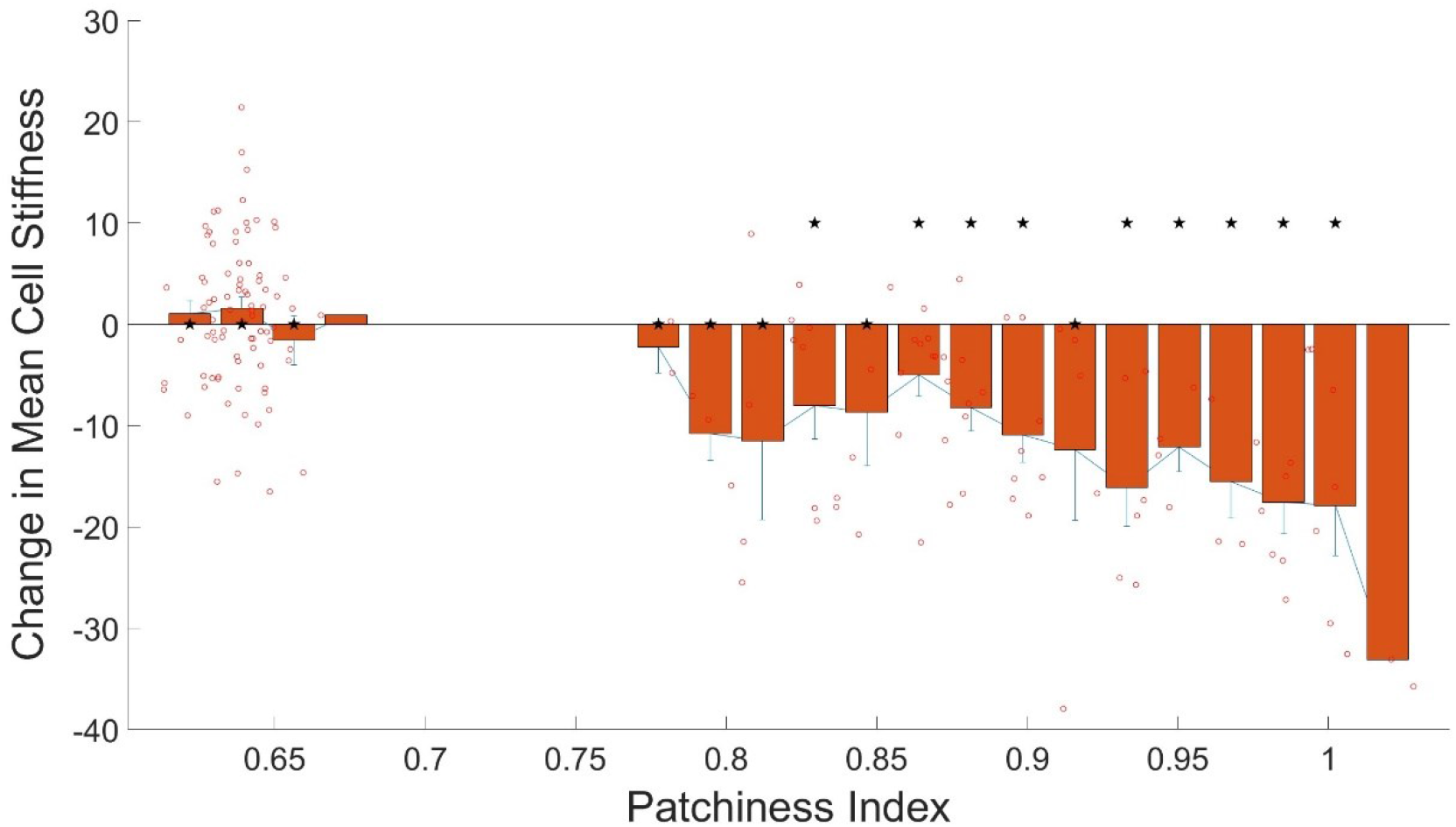
Change in mean stiffness of the cells against c value (ratio of local to global diversity). Window size of 4×4×4 cell lengths (40 µm × 40 µm × 40 µm) used to calculate the patchiness index using equation 5. Each open red circle represents the change in mean cell stiffness over 20 cell death and division cycles for a single simulation, plotted against the patchiness index of the starting initial tissue configuration for that simulation run. The error bars are standard errors of the mean. Stars above the bar plots indicate p values < 0.05 for a single population ttest.

## Discussion

Here we have used a purely theoretical setup to study the evolution of a mechanically diverse population of cancer cells within a densely packed 3D tissue environment. We have ignored the presence of nutrient gradients, extracellular matrix, or significantly different cell types, concentrating only epithelial or near epithelial cell types. Cell-cell interactions are assumed to be purely mechanical and cell fate decisions are also driven primarily by mechanosensitive feedback based on empirical experimental observations. While all these amount to a gross simplification of a complex tissue system, such simplified models have previously been employed to understand and describe processes related to tissue dynamics (44,45), tissue growth (46), tumor incidence (3), tumor growth and metastatic invasion (47–49). Additionally, the small length scale of the tissue system simulated in this case allows for the assumption of a dense collection of tightly packed cells without the inclusion of an extracellular matrix, nutrient gradients, or vasculature. While additional complexity can be added to the model, we leave these for future extensions of the model.

Within the confines of the limitations described above, we use our model to understand how heterogeneity in the mechanical properties of tumor cells, specifically, cell stiffness, as observed and quantified by experimental tools, may be used to predict the occurrence of tumor growth and a malignant shift in a seemingly healthy population of cells. We find that heterogeneity at the global level, but homogeneity at a local level, quantified by the patchiness of cell distributions within the tissue system, predicts an increasing likelihood of tumor occurrence and an overall shift towards malignant populations within an initially health tissue. It is interesting to note that overall high heterogeneity, by itself is not detrimental to tissue fate, so long as the population remains well mixed. However, local clustering of similar populations, which may occur due to stochastic cell division events, or due to migration-based segregation (28,29,50), tips the balance to drive malignant transformation in highly heterogenous tissue systems.

We have restricted this study to the heterogeneity in the cell mechanotype, as defined by cell stiffness, since this is one of the key mechanical properties known to distinguish cancerous vs normal cell populations and can also help grade cancer cell populations based on their aggressiveness (51–54). Additionally, there are tools being developed that can measure and map cellular mechanical stiffness in situ within tissue samples (55,56). However, other cellular mechanical properties such as cell adhesion strength, cell contractility and cell nuclear deformability also show similar heterogeneity levels within genetically identical cell populations (57–59). Both cell adhesion strength and cell contractility have similar effects to that of cell stiffness on cell organization, shape, and size within vertex-based cell-cell interaction models (60), and we believe will lead to similar observations as those presented. There are few models currently incorporating cell nuclear stiffness when considering cell-cell interactions and tissue dynamics, and this is something that needs to be worked on in the future. Lastly, these mechanical differences between cells arise from biochemical differences driven by differences in gene expression profiles within a cell population. Thus, cell phenotypic differences may directly correlate to cell mechanotype differences, and the analysis of tissue patchiness may be extended to any spatial characterization of cells within a tissue environment. Indeed, with invention of accurate single cell gene expression analysis and tracking tools, as well as large scale spatial phenotypical and mechanical profiling of cells in-situ (61,62), it might be possible to experimentally characterize a tissue based on its patchiness. Based on the results presented here, we propose that tissue patchiness can provide key diagnostic and prognostic insights for malignant growth in health tissues and benign tumors.

Heterogeneity is a norm in biology and cannot be done away with. This is true across scales and species. On the other hand, limiting population de-mixing and clustering into local niches may be a potential strategy that can be employed to avoid tipping of the scales and allowing one sub-species to dominate. Patchiness may be avoided by limiting factors that promote proliferation of only certain sub-populations, or factors that limit the mobility of cells within certain regions of the tissue. Biological events and biochemical or biomechanical factors that potentially increase or decrease patchiness within normal tissues or tumors, and their relation to actual tumor growth and malignancy needs to be further investigated and discussed. However, we strongly believe that the idea of patchiness might be just as applicable to cancer ecology as it is to environmental ecology and population dynamics.

## Conclusion

Here we present a model that focuses on cell interactions and mechanoreciprocity as drivers of tissue dynamics. We use this model to understand how global and local heterogeneity in cellular mechanics across a dense population of cells may lead to an incidence of malignant tumor growth defined by an increase in the population of cancer-like soft cells and decrease in the population of normal, stiff cells. Based on our results, we find that tissue patchiness as defined by the ratio of global to local heterogeneity may be an excellent metric to predict malignant transformation in healthy tissues or benign tumors. While limited by the purely theoretical nature of this study and a sole focus on cell mechanics, the potential for such a metric is extremely appealing for early diagnosis and intervention in cancer patients as well as prognosis in patients with benign tumors. Additionally, the model is not only limited to studying the effects of cellular mechanical properties on tumorigenesis in a healthy tissue, but it can also serve in studying a broader scope of tissue mechanics in processes like wound healing and aging. Our model also provides a versatile platform that can be built upon, with the ability to study the effects other mechanical properties that have been linked to cancerous behavior or potential interventional strategies focused on manipulating the ecological landscape of tissues and tumors.

## Acknowledgements

P.K. acknowledges funding for this project by the Army Research Office grant W911NF-17-1-0413.

## References

1. Suresh S. Biomechanics and biophysics of cancer cells☆. Acta Biomater. 2007 Jul;3(4):413–38.

2. Katira P, Bonnecaze RT, Zaman MH. Modeling the Mechanics of Cancer: Effect of Changes in Cellular and Extra-Cellular Mechanical Properties. Front Oncol [Internet]. 2013 [cited 2022 May 26];3. Available from: http://journal.frontiersin.org/article/10.3389/fonc.2013.00145/abstract

3. Katira P, Zaman MH, Bonnecaze RT. How Changes in Cell Mechanical Properties Induce Cancerous Behavior. Phys Rev Lett. 2012 Jan 11;108(2):028103.

4. Kim TH, Rowat AC, Sloan EK. Neural regulation of cancer: from mechanobiology to inflammation. Clin Transl Immunol. 2016 May 13;5(5):e78.

5. Jaalouk DE, Lammerding J. Mechanotransduction gone awry. Nat Rev Mol Cell Biol. 2009 Jan;10(1):63–73.

6. Hanahan D, Weinberg RA. Hallmarks of Cancer: The Next Generation. Cell. 2011 Mar;144(5):646–74.

7. Levental KR, Yu H, Kass L, Lakins JN, Egeblad M, Erler JT, et al. Matrix Crosslinking Forces Tumor Progression by Enhancing Integrin Signaling. Cell. 2009 Nov;139(5):891–906.

8. Kraning-Rush CM, Califano JP, Reinhart-King CA. Cellular Traction Stresses Increase with Increasing Metastatic Potential. Laird EG, editor. PLoS ONE. 2012 Feb 28;7(2):e32572.

9. Omidvar R, Tafazzoli-shadpour M, Shokrgozar MA, Rostami M. Atomic force microscope-based single cell force spectroscopy of breast cancer cell lines: An approach for evaluating cellular invasion. J Biomech. 2014 Oct;47(13):3373–9.

10. Li QS, Lee GYH, Ong CN, Lim CT. AFM indentation study of breast cancer cells. Biochem Biophys Res Commun. 2008 Oct;374(4):609–13.

11. Luo Q, Kuang D, Zhang B, Song G. Cell stiffness determined by atomic force microscopy and its correlation with cell motility. Biochim Biophys Acta BBA - Gen Subj. 2016 Sep;1860(9):1953–60.

12. Alibert C, Goud B, Manneville JB. Are cancer cells really softer than normal cells?: Mechanics of cancer cells. Biol Cell. 2017 May;109(5):167–89.

13. Fritsch A, Höckel M, Kiessling T, Nnetu KD, Wetzel F, Zink M, et al. Are biomechanical changes necessary for tumour progression? Nat Phys. 2010 Oct;6(10):730–2.

14. Xu W, Mezencev R, Kim B, Wang L, McDonald J, Sulchek T. Cell Stiffness Is a Biomarker of the Metastatic Potential of Ovarian Cancer Cells. Batra SK, editor. PLoS ONE. 2012 Oct 4;7(10):e46609.

15. Swaminathan V, Mythreye K, O’Brien ET, Berchuck A, Blobe GC, Superfine R. Mechanical Stiffness Grades Metastatic Potential in Patient Tumor Cells and in Cancer Cell Lines. Cancer Res. 2011 Aug 1;71(15):5075–80.

16. Lekka M, Gil D, Pogoda K, Dulińska-Litewka J, Jach R, Gostek J, et al. Cancer cell detection in tissue sections using AFM. Arch Biochem Biophys. 2012 Feb;518(2):151–6.

17. Cross SE, Jin YS, Rao J, Gimzewski JK. Nanomechanical analysis of cells from cancer patients. Nat Nanotechnol. 2007 Dec;2(12):780–3.

18. Nikkhah M, Strobl JS, Schmelz EM, Agah M. Evaluation of the influence of growth medium composition on cell elasticity. J Biomech. 2011 Feb;44(4):762–6.

19. Lekka M, Laidler P, Gil D, Lekki J, Stachura Z, Hrynkiewicz AZ. Elasticity of normal and cancerous human bladder cells studied by scanning force microscopy. Eur Biophys J. 1999 May 25;28(4):312–6.

20. Ramos JR, Pabijan J, Garcia R, Lekka M. The softening of human bladder cancer cells happens at an early stage of the malignancy process. Beilstein J Nanotechnol. 2014 Apr 10;5:447–57.

21. Plodinec M, Loparic M, Monnier CA, Obermann EC, Zanetti-Dallenbach R, Oertle P, et al. The nanomechanical signature of breast cancer. Nat Nanotechnol. 2012 Oct 21;7(11):757–65.

22. Fuhs T, Wetzel F, Fritsch AW, Li X, Stange R, Pawlizak S, et al. Rigid tumours contain soft cancer cells. Nat Phys. 2022 Dec;18(12):1510–9.

23. Gottheil P, Lippoldt J, Grosser S, Renner F, Saibah M, Tschodu D, et al. State of Cell Unjamming Correlates with Distant Metastasis in Cancer Patients. Phys Rev X. 2023 Jul 10;13(3):031003.

24. Efremov YuM, Lomakina ME, Bagrov DV, Makhnovskiy PI, Alexandrova AY, Kirpichnikov MP, et al. Mechanical properties of fibroblasts depend on level of cancer transformation. Biochim Biophys Acta BBA - Mol Cell Res. 2014 May;1843(5):1013–9.

25. Altrock PM, Liu LL, Michor F. The mathematics of cancer: integrating quantitative models. Nat Rev Cancer. 2015 Dec;15(12):730–45.

26. Macklin P, McDougall S, Anderson ARA, Chaplain MAJ, Cristini V, Lowengrub J. Multiscale modelling and nonlinear simulation of vascular tumour growth. J Math Biol. 2009 Apr;58(4–5):765–98.

27. Lowengrub JS, Frieboes HB, Jin F, Chuang YL, Li X, Macklin P, et al. Nonlinear modelling of cancer: bridging the gap between cells and tumours. Nonlinearity. 2010 Jan 1;23(1):R1–91.

28. Reddy GA, Katira P. Differences in cell death and division rules can alter tissue rigidity and fluidization. Soft Matter. 2022;18(19):3713–24.

29. Heine P, Lippoldt J, Reddy GA, Katira P, Käs JA. Anomalous cell sorting behavior in mixed monolayers discloses hidden system complexities. New J Phys. 2021 Apr 1;23(4):043034.

30. Chen CS, Mrksich M, Huang S, Whitesides GM, Ingber DE. Geometric Control of Cell Life and Death. Science. 1997 May 30;276(5317):1425–8.

31. Dike LE, Chen CS, Mrksich M, Tien J, Whitesides GM, Ingber DE. Geometric control of switching between growth, apoptosis, and differentiation during angiogenesis using micropatterned substrates. Vitro Cell Dev Biol - Anim. 1999 Sep;35(8):441–8.

32. Jonas O, Mierke CT, Käs JA. Invasive cancer cell lines exhibit biomechanical properties that are distinct from their noninvasive counterparts. Soft Matter. 2011;7(24):11488.

33. Atia L, Bi D, Sharma Y, Mitchel JA, Gweon B, A. Koehler S, et al. Geometric constraints during epithelial jamming. Nat Phys. 2018 Jun;14(6):613–20.

34. Köppen M, Fernández BG, Carvalho L, Jacinto A, Heisenberg CP. Coordinated cell-shape changes control epithelial movement in zebrafish and Drosophila. Development. 2006 Jul 15;133(14):2671–81.

35. Rosenblatt J, Raff MC, Cramer LP. An epithelial cell destined for apoptosis signals its neighbors to extrude it by an actin- and myosin-dependent mechanism. Curr Biol. 2001 Nov;11(23):1847–57.

36. Dolznig H, Grebien F, Sauer T, Beug H, Müllner EW. Evidence for a size-sensing mechanism in animal cells. Nat Cell Biol. 2004 Sep;6(9):899–905.

37. Streichan SJ, Hoerner CR, Schneidt T, Holzer D, Hufnagel L. Spatial constraints control cell proliferation in tissues. Proc Natl Acad Sci. 2014 Apr 15;111(15):5586–91.

38. Chaffey N. Alberts, B., Johnson, A., Lewis, J., Raff, M., Roberts, K. and Walter, P. Molecular biology of the cell. 4th edn. Ann Bot. 2003 Feb 1;91(3):401–401.

39. Caswell H, Cohen JE. Communities in Patchy Environments: A Model of Disturbance, Competition, and Heterogeneity. In: Kolasa J, Pickett STA, editors. Ecological Heterogeneity [Internet]. New York, NY: Springer New York; 1991 [cited 2023 Jul 30]. p. 97–122. (Billings WD, Golley F, Lange OL, Olson JS, Remmert H, editors. Ecological Studies; vol. 86). Available from: http://link.springer.com/10.1007/978-1-4612-3062-5_6

40. Marquet PA, Fortin MJ, Pineda J, Wallin DO, Clark J, Wu Y, et al. Ecological and Evolutionary Consequences of Patchiness: A Marine-Terrestrial Perspective. In: Levin SA, Powell TM, Steele JW, editors. Patch Dynamics [Internet]. Berlin, Heidelberg: Springer Berlin Heidelberg; 1993 [cited 2023 Jul 30]. p. 277–304. (Levin SA, editor. Lecture Notes in Biomathematics; vol. 96). Available from: http://link.springer.com/10.1007/978-3-642-50155-5_19

41. Horne JK, Schneider DC. Spatial Variance in Ecology. Oikos. 1995 Oct;74(1):18.

42. Patch dynamics | Ecological Succession, Species Interactions & Landscape Ecology | Britannica [Internet]. [cited 2023 Jul 31]. Available from: https://www.britannica.com/science/patch-dynamics

43. Helmlinger G, Netti PA, Lichtenbeld HC, Melder RJ, Jain RK. Solid stress inhibits the growth of multicellular tumor spheroids. Nat Biotechnol. 1997 Aug 1;15(8):778–83.

44. Park JA, Kim JH, Bi D, Mitchel JA, Qazvini NT, Tantisira K, et al. Unjamming and cell shape in the asthmatic airway epithelium. Nat Mater. 2015 Oct;14(10):1040–8.

45. Fletcher AG, Osterfield M, Baker RE, Shvartsman SY. Vertex Models of Epithelial Morphogenesis. Biophys J. 2014 Jun 3;106(11):2291–304.

46. Lin SZ, Li B, Feng XQ. A dynamic cellular vertex model of growing epithelial tissues. Acta Mech Sin. 2017 Apr 1;33(2):250–9.

47. Bi D, Lopez JH, Schwarz JM, Manning ML. Energy barriers and cell migration in densely packed tissues. Soft Matter. 2014;10(12):1885.

48. Grosser S, Lippoldt J, Oswald L, Merkel M, Sussman DM, Renner F, et al. Cell and Nucleus Shape as an Indicator of Tissue Fluidity in Carcinoma. Phys Rev X. 2021 Feb 17;11(1):011033.

49. Alt S, Ganguly P, Salbreux G. Vertex models: from cell mechanics to tissue morphogenesis. Philos Trans R Soc B Biol Sci. 2017 Mar 27;372(1720):20150520.

50. Brodland GW. Computational modeling of cell sorting, tissue engulfment, and related phenomena: A review. Appl Mech Rev. 2004;57(1):47.

51. Xu W, Mezencev R, Kim B, Wang L, McDonald J, Sulchek T. Cell Stiffness Is a Biomarker of the Metastatic Potential of Ovarian Cancer Cells. Batra SK, editor. PLoS ONE. 2012 Oct 4;7(10):e46609.

52. Swaminathan V, Mythreye K, O’Brien ET, Berchuck A, Blobe GC, Superfine R. Mechanical Stiffness Grades Metastatic Potential in Patient Tumor Cells and in Cancer Cell Lines. Cancer Res. 2011 Aug 1;71(15):5075–80.

53. Gill NK, Ly C, Nyberg KD, Lee L, Qi D, Tofig B, et al. A scalable filtration method for high throughput screening based on cell deformability. Lab Chip. 2019 Jan 15;19(2):343–57.

54. Otto O, Rosendahl P, Mietke A, Golfier S, Herold C, Klaue D, et al. Real-time deformability cytometry: on-the-fly cell mechanical phenotyping. Nat Methods. 2015 Mar;12(3):199–202.

55. Morr AS, Nowicki M, Bertalan G, Vieira Silva R, Infante Duarte C, Koch SP, et al. Mechanical properties of murine hippocampal subregions investigated by atomic force microscopy and in vivo magnetic resonance elastography. Sci Rep. 2022 Oct 6;12(1):16723.

56. Plodinec M, Lim RYH. Nanomechanical Characterization of Living Mammary Tissues by Atomic Force Microscopy. In: Vivanco M del M, editor. Mammary Stem Cells: Methods and Protocols [Internet]. New York, NY: Springer; 2015 [cited 2023 Jul 31]. p. 231–46. (Methods in Molecular Biology). Available from: https://doi.org/10.1007/978-1-4939-2519-3_14

57. Beri P, Popravko A, Yeoman B, Kumar A, Chen K, Hodzic E, et al. Cell Adhesiveness Serves as a Biophysical Marker for Metastatic Potential. Cancer Res. 2020 Feb 14;80(4):901–11.

58. Bordeleau F, Reinhart-King CA. Tuning cell migration: contractility as an integrator of intracellular signals from multiple cues. F1000Research [Internet]. 2016 Jul 26 [cited 2016 Sep 25];5. Available from: http://www.ncbi.nlm.nih.gov/pmc/articles/PMC4962296/

59. Khan ZS, Santos JM, Hussain F. Aggressive prostate cancer cell nuclei have reduced stiffness. Biomicrofluidics. 2018 Jan 2;12(1):014102.

60. Manning ML, Foty RA, Steinberg MS, Schoetz EM. Coaction of intercellular adhesion and cortical tension specifies tissue surface tension. Proc Natl Acad Sci. 2010 Jul 13;107(28):12517–22.

61. Bouwman BAM, Crosetto N, Bienko M. The era of 3D and spatial genomics. Trends Genet. 2022 Oct 1;38(10):1062–75.

62. Zhao T, Chiang ZD, Morriss JW, LaFave LM, Murray EM, Del Priore I, et al. Spatial genomics enables multi-modal study of clonal heterogeneity in tissues. Nature. 2022 Jan;601(7891):85–91.

63. Nyberg KD, Scott MB, Bruce SL, Gopinath AB, Bikos D, Mason TG, et al. The physical origins of transit time measurements for rapid, single cell mechanotyping. Lab Chip. 2016;16(17):3330–9.

64. Xu X, Li Z, Cai L, Calve S, Neu CP. Mapping the Nonreciprocal Micromechanics of Individual Cells and the Surrounding Matrix Within Living Tissues. Sci Rep. 2016 Jul;6(1):24272.

65. Koch TM, Münster S, Bonakdar N, Butler JP, Fabry B. 3D Traction Forces in Cancer Cell Invasion. Mellor H, editor. PLoS ONE. 2012 Mar 30;7(3):e33476.

66. Phillip JM, Aifuwa I, Walston J, Wirtz D. The Mechanobiology of Aging. Annu Rev Biomed Eng. 2015 Dec 7;17(1):113–41.

