## Supplementary Information for "“Patchiness” in Mechanical Stiffness across a Tumor as an Early-Stage Marker for Malignancy"

**Table 1 Parameters and Descriptions.**

| Parameter | Value | Description | Reference |
|---|---|---|---|
| $V_0$ | 1000 µm$^3$ | Volume of spherical cell | [1] |
| E | Varies. Taken from a log-normal or normal distribution with a mean value of 500 Pa and particular variations. | Cell actin cortex elastic modulus | [2–6] |
| Y | 1 µm | Actin cortex thickness | [7,8] |
| Σ | 200 µm$^{-2}$ | Bond density between all cells | [9] |
| γ | 20$k_bT$ | Bond energy of intercellular bonds | [10,11] |
| $\pi_0$ | 100 Pa | Osmotic pressure | [12] |

**Table 2. Different initially healthy tissue systems simulated.**

| Type of Simulation | Spatial Arrangement | Distribution | Variation |
|---|---|---|---|
| 1 | 8 seed cells | Log-Normal | Mode: 425 Pa |
| 2 | 27 seed cells | Log-Normal | Mode: 425 Pa |
| 3 | 64 seed cells | Log-Normal | Mode: 425 Pa |
| 4 | Non-Clustered | Log-Normal | Mode: 425 Pa |
| 5 | 8 seed cells | Log-Normal | Mode: 435 Pa |
| 6 | 27 seed cells | Log-Normal | Mode: 435 Pa |
| 7 | 64 seed cells | Log-Normal | Mode: 435 Pa |
| 8 | Non-Clustered | Log-Normal | Mode: 435 Pa |
| 9 | 8 seed cells | Log-Normal | Mode: 445 Pa |
| 10 | 27 seed cells | Log-Normal | Mode: 445 Pa |
| 11 | 64 seed cells | Log-Normal | Mode: 445 Pa |
| 12 | Non-Clustered | Log-Normal | Mode: 445 Pa |
| 13 | 8 seed cells | Log-Normal | Mode: 455 Pa |
| 14 | 27 seed cells | Log-Normal | Mode: 455 Pa |
| 15 | 64 seed cells | Log-Normal | Mode: 455 Pa |
| 16 | Non-Clustered | Log-Normal | Mode: 455 Pa |
| 17 | 8 seed cells | Log-Normal | Mode: 465 Pa |
| 18 | 27 seed cells | Log-Normal | Mode: 465 Pa |
| 19 | 64 seed cells | Log-Normal | Mode: 465 Pa |
| 20 | Non-Clustered | Log-Normal | Mode: 465 Pa |

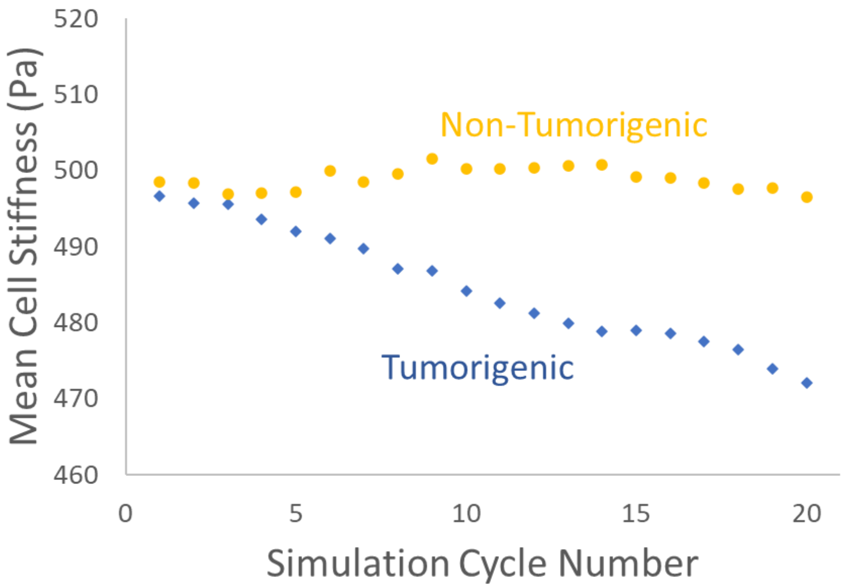


Figure 1. In tissues where tumorigenesis occurs, there is a constant decrease in the mean value of the cell stiffness distribution over time. In tissues where tumorigenesis does not occur, the mean value of the distribution remains constant over time or may go up. The figure shows two sample runs, one with intermediate clustering (27 seed cells) and low initial variance in cell stiffness (Mode 465, Mean 500) (orange circles) and another with high cluster (8 seed cells) and high initial variance in cell stiffness (Mode 425, Mean 500) (blue diamonds).

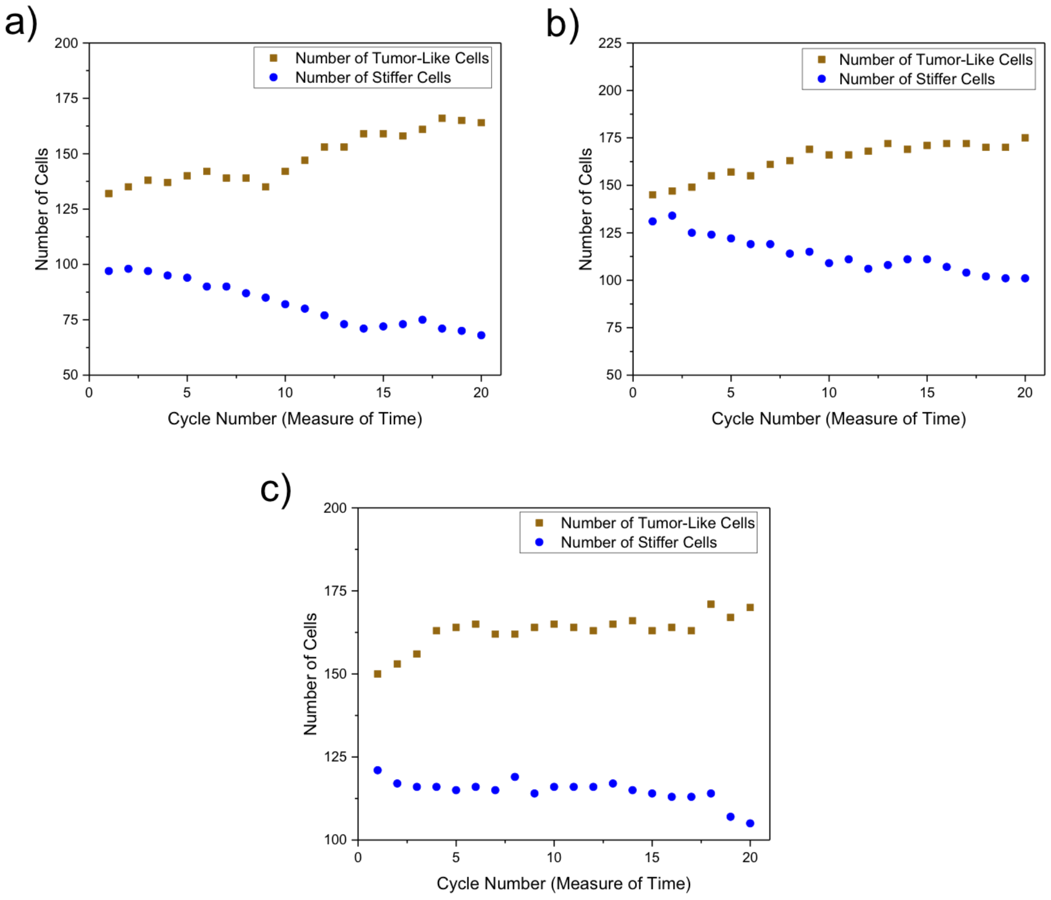


Figure 2. Negative shifts in the mean of the distribution of at least 15 Pa in the tissue are observed with increases in amount of tumor-like cells and decreases in the amount of stiffer cells. a) a spatially heterogeneous tissue with 8 seed cells and a log-normal mode of 445 Pa, b) a spatially heterogeneous tissue with 8 seed cells and a log-normal mode of 425 Pa, c) a spatially heterogeneous with 27 seed cells and a log-normal mode of 435 Pa.

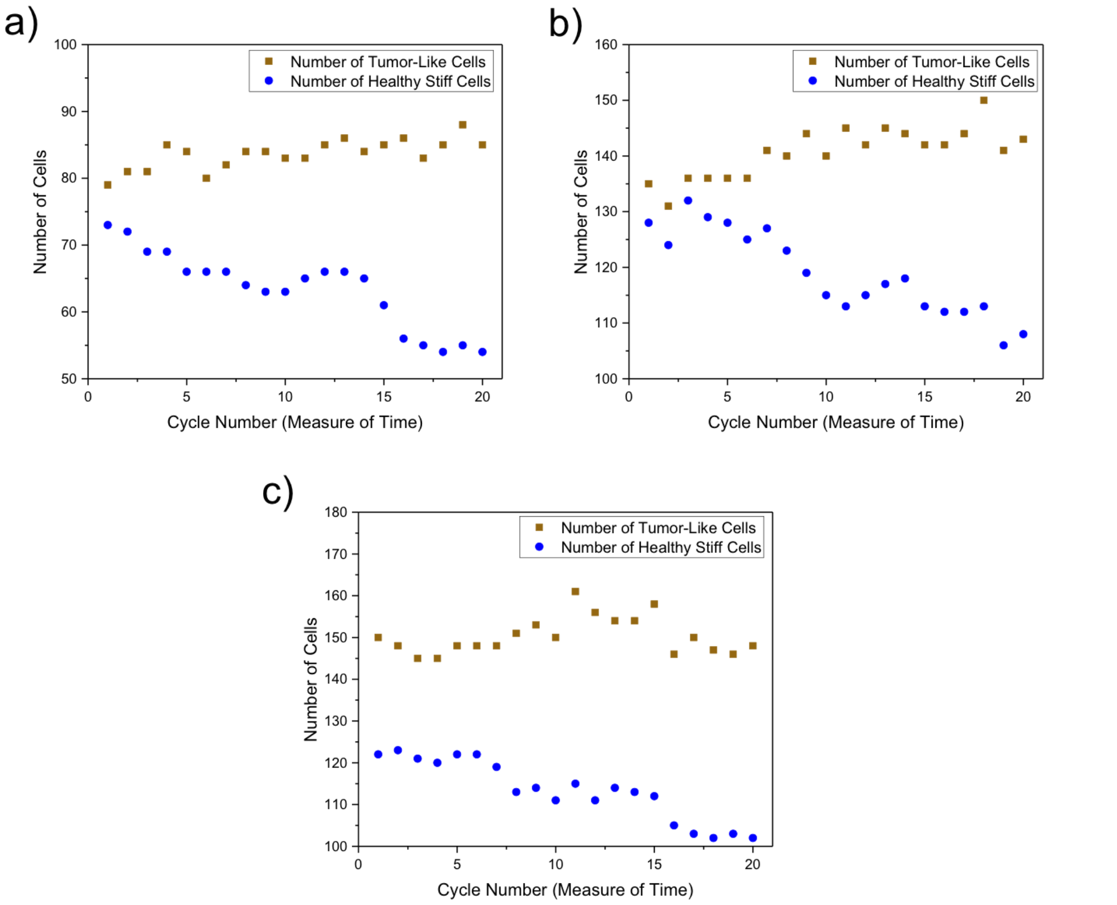


Figure 3. Negative shifts in the mean of the distribution of at least 15 Pa in the tissue are observed with a relatively constant amount of tumor-like cells and a substantial decrease in the amount of stiffer cells. a) a spatially heterogeneous tissue with 8 seed cells and a log-normal mode of 465 Pa, b) a spatially heterogeneous tissue with 27 seed cells and a log-normal mode of 435 Pa, c) a spatially heterogeneous with 27 seed cells and a log-normal mode of 435 Pa.

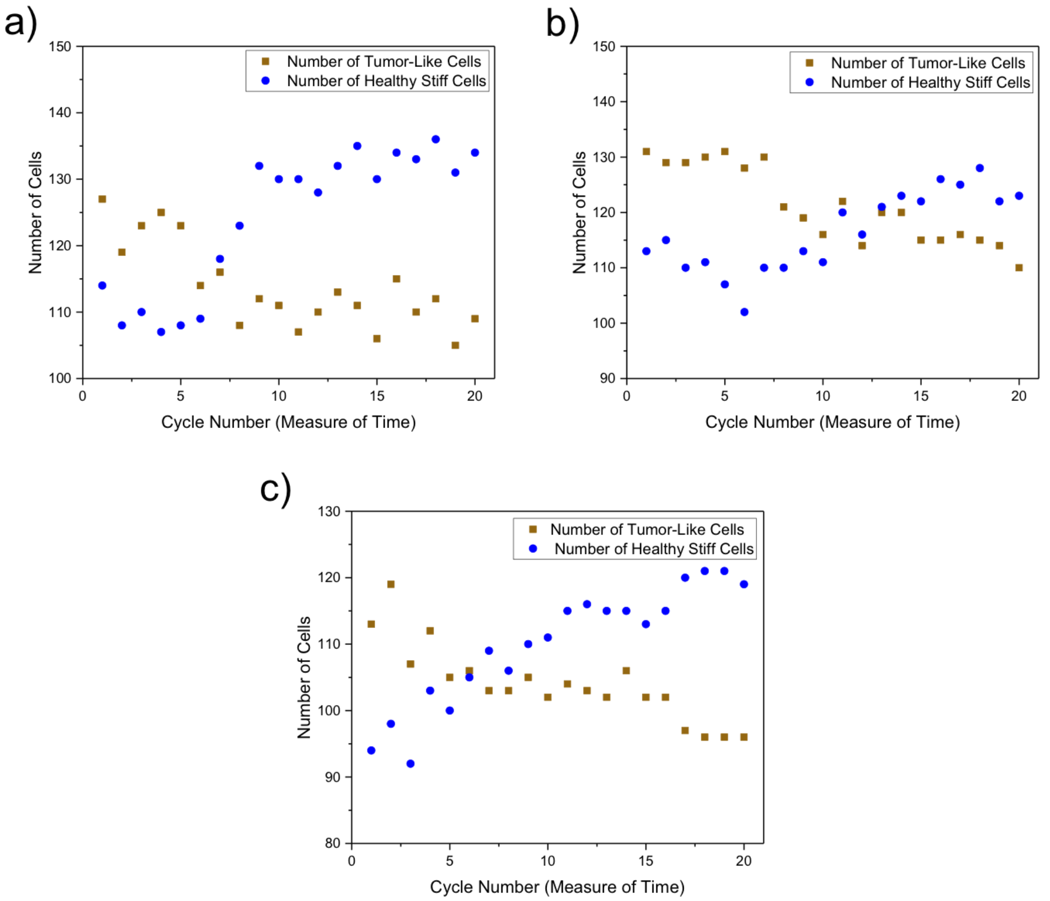


Figure 4. Positive shifts in the mean of the distribution of at least 15 Pa in the tissue are observed with an increase of stiffer cells and a gradual decrease in the amount of tumor-like cells. a) a spatially homogeneous tissue with a log-normal mode of 445 Pa, b) a spatially homogeneous tissue with a log-normal mode of 445 Pa, c) a spatially homogeneous tissue with a log-normal mode of 455 Pa.

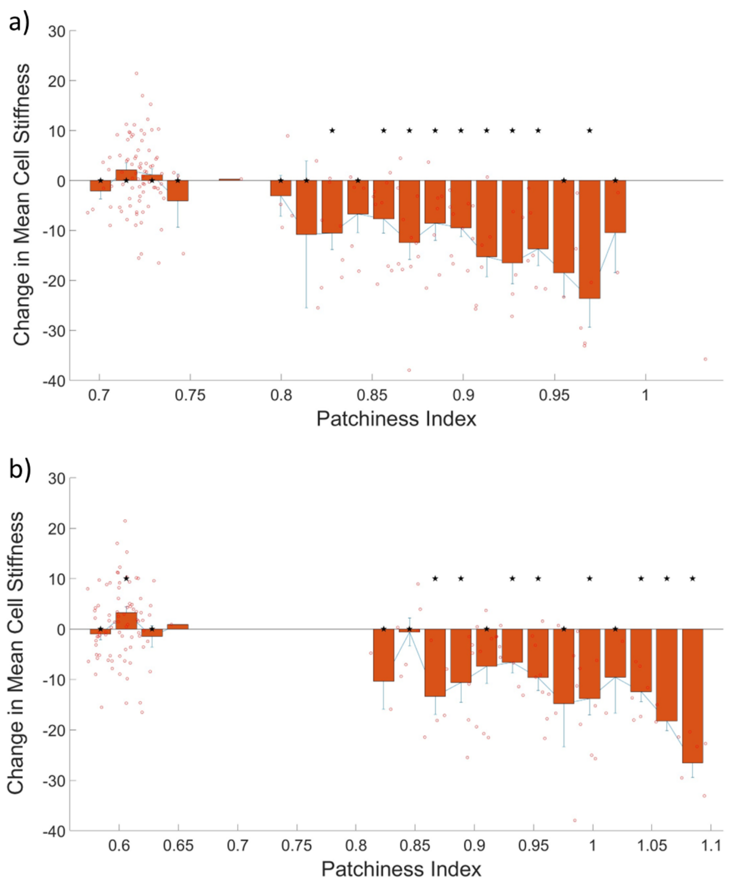


Figure 5. Change in mean stiffness of the cells against c value (ratio of local to global diversity). a) Window size of 3x3x3 cell lengths (30 µm x 30 µm x 30 µm) and b) Window size of 5x5x5 cell lengths (50 µm x 50 µm x 50 µm) used to calculate the patchiness index using equation 5. Each open red circle represents the change in mean cell stiffness over 20 cell death and division cycles for a single simulation, plotted against the patchiness index of the starting initial tissue configuration for that simulation run. The error bars are standard errors of the mean. Stars above the bar plots indicate p values < 0.05 for a single population ttest.

1. Katira, P., Zaman, M. H. & Bonnecaze, R. T. How changes in cell mechanical properties induce cancerous behavior. *Physical Review Letters* **108**, 1–5 (2012).
2. Jonas, O., Mierke, C. T. & Käs, J. A. Invasive cancer cell lines exhibit biomechanical properties that are distinct from their noninvasive counterparts. *Soft Matter* **7**, 11488 (2011).
3. Lekka, M. *et al.* Cancer cell detection in tissue sections using AFM. *Archives of Biochemistry and Biophysics* **518**, 151–156 (2012).
4. Sun, J. *et al.* Biomechanical profile of cancer stem-like cells derived from MHCC97H cell lines. *Journal of Biomechanics* **49**, 45–52 (2016).
5. Efremov, Y. M. *et al.* Mechanical properties of fibroblasts depend on level of cancer transformation. *Biochimica et Biophysica Acta - Molecular Cell Research* **1843**, 1013–1019 (2014).
6. Xu, W. *et al.* Cell Stiffness Is a Biomarker of the Metastatic Potential of Ovarian Cancer Cells. *PLoS ONE* **7**, (2012).
7. Wottawah, F. *et al.* Optical rheology of biological cells. *Physical Review Letters* **94**, 1–4 (2005).
8. Ananthakrishnan, R. *et al.* Quantifying the contribution of actin networks to the elastic strength of fibroblasts. *Journal of Theoretical Biology* **242**, 502–516 (2006).
9. Li, Z. *et al.* Spatially resolved quantification of E-cadherin on target hES cells. *Journal of Physical Chemistry B* **114**, 2894–2900 (2010).
10. Zhu, B. *et al.* Functional analysis of the structural basis of homophilic cadherin adhesion. *Biophysical Journal* **84**, 4033–4042 (2003).

11. Byers, S. W., Sommers, C. L., Hoxter, B., Mercurio, a M. & Tozeren, a. Role of E-cadherin in the response of tumor cell aggregates to lymphatic, venous and arterial flow: measurement of cell-cell adhesion strength. *Journal of cell science* **108 ( Pt 5**, 2053–2064 (1995).

12. Charras, G. T., Coughlin, M., Mitchison, T. J. & Mahadevan, L. Life and Times of a Cellular Bleb. *Biophysical Journal* **94**, 1836–1853 (2008).
